# A multiscale model of the mammalian liver circadian clock supports synchronization of autonomous oscillations by intercellular communication

**DOI:** 10.1101/2024.02.15.580517

**Authors:** Daniel Marri, Omar Kana, David Filipovic, James P. Sluka, Shengnan Liu, Qiang Zhang, Sudin Bhattacharya

## Abstract

Expression of core circadian clock genes in hepatocytes across the liver lobule is temporally synchronized despite cell-autonomous oscillations in gene expression. This spatial synchronization has been attributed to an unknown intercellular coupling mechanism. Here we have developed multicellular computational models of the murine liver lobule with and without intercellular coupling to investigate the role of synchronization in circadian gene expression. Our models demonstrated that intercellular coupling was needed to generate sustained circadian oscillations with a near 24-hour period. Without coupling the simulated period was variable within the 21-28-hour range. Further model analysis revealed that a robust near-24-hour oscillation period can be generated with a wide range of circadian protein degradation rates. In contrast, only a small window of circadian gene transcription rates was able to generate realistic oscillatory periods. The coupled model accurately captured the temporal dynamics of circadian genes derived from single-nuclei transcriptomic data. Overall, this study provides novel insights into the mammalian hepatic circadian clock through modeling of spatial and temporal gene expression patterns and data-driven analysis.

## Introduction

Species ranging from cyanobacteria to mammals have an inbuilt biological clock capable of generating sustained oscillations with a period close to 24 hr^1,2^. These circadian oscillations are endogenous and persist in the absence of environmental timing cues. The suprachiasmatic nucleus (SCN) in the brain serves as the master pacemaker of the circadian clock and synchronizes cellular, tissue, and systemic rhythms to regulate varied biological processes^3,4^ like maintenance of body temperature, glucose metabolism, sleep-wake cycle, hormone regulation, and bone formation^5–7^. Disruption of the circadian cycle by environmental stimuli has been associated with cardiovascular disease, diabetes, bipolar disorder, obesity, and cancer^8–11^. A network of transcriptional and translational and post translation feedback loops underlies the biological clock that generates circadian rhythms in the SCN and peripheral tissues like the liver^12,13^. The core of this network is a set of transcriptional activators: circadian locomotor output cycles kaput (*Clock*), Neuronal PAS domain protein 2 (*Npas2*), brain and muscle ARNT Like 1 (*Bmal1*), retinoic acid-related orphan receptor (*Rora, Rorb, Rorc*); and repressors: the period genes (*Per1, Per2, Per3*), the cryptochrome genes (*Cry1, Cry2*) and reverb-clear orphan receptors (*Reverbα, Reverbβ*)^3,14^. The master regulatory heterodimer CLOCK-BMAL1 (or NPAS2-BMAL1) binds to the E-box DNA motif in regulatory regions of the rhythmic genes *Per, Cry, Ror, and Reverb* to activate their transcription. The PER and CRY cytosolic proteins form another heterodimer complex which is translocated into the nucleus to inhibit transcription of their own genes in an autoregulatory negative feedback loop. The heterodimer also suppresses the transcription of Ror and Reverb genes. ROR and REV-ERB proteins compete for ROR regulatory element (RRE) binding sites in the promoter region of *Bmal1* to regulate its transcription; with ROR serving as an activator and REV-ERB as a repressor^13,15^.

The liver is a peripheral oscillator acting as an essential site for anabolic and catabolic processing of lipids and amino acids^16^. The mammalian liver is spatially organized into functional sub-units called “lobules” composed primarily of hepatocytes, the cell type making up the liver parenchyma. Hepatocytes lie along a lobular axis extending from a portal vein to a central vein and thus can be categorized according to their proximity to either the central or portal vein (Fig 1). Gene expression and resulting metabolic functions show a spatial gradation along the portal-to-central axis of the lobule, with gluconeogenesis and β-oxidation enriched at the portal end, and glycolysis and lipogenesis enriched towards the central region^17^. In contrast, a recent single-cell gene expression study showed that the core circadian clock genes are expressed in a non-zonated manner, i.e., there is no significant difference in their expression in hepatocytes across the portal-to-central axis^18^. This observation has been attributed to a coupling mechanism that synchronizes the gene expression among cells across the portal-to-central axis to produce coordinated circadian oscilaltions^19–21^. While the coupling between cells in the SCN is understood to be partly achieved by neurotransmitters,^22^ the precise mechanism responsible for synchronized oscillations in liver gene expression is not fully understood, though transforming growth factor–b (TGF-b) has been proposed as a putative coupling factor^21,23,24^.

**Figure 1:**
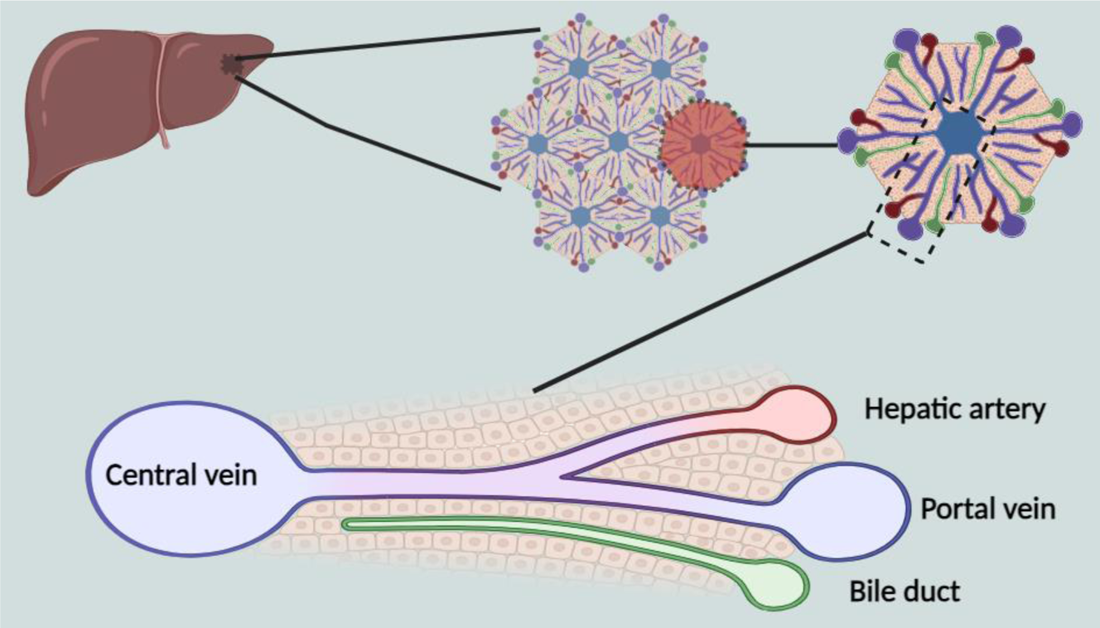
Schematic of a cross-section through the liver lobule. The fundamental structural unit of the mammalian liver is the hepatic lobule, predominantly made up of hepatocytes extending in layers from the portal triad to the central vein (generated using https://biorender.com/).

In this study, we present a spatiotemporal multicellular mathematical model of the mammalian liver circadian clock regulatory network.^25–28^ We developed two sets of models of the mouse hepatic clock: 1) Model 1, with no communication among the cellular oscillators leading to non-synchronized gene expression in hepatocytes across the central to portal axis of the lobule; and 2) Model 2, with communication among cells leading to synchronized gene expression across the lobular axis. We found a positive relation between the amplitudes of the observables and their transcription rate parameters, and a negative relation between the amplitudes and their respective degradation rates. Finally, we estimated the transcription and degradation rate parameters in our model from single-cell RNA-Seq data^25^. This analysis yielded a high correlation between simulated and experimental data with an *R^2^* > 0.9. In summary, we computationally analyzed asynchronous and synchronous spatial and temporal circadian oscillations in the mammalian liver, as well as the dependence of the model observables on their respective transcription and degradation rate parameters.

## Results

### Circadian clock mechanism – model design

To develop our model, we identified and compiled from literature the key regulatory interactions in the mammalian clock gene network^26,27^. Six known protein-coding clock genes CLOCK< BMAL1, *PER, CRY, ROR, and REV-ERB* were used to develop the model. For simplicity, we did not distinguish between multiple isoforms of these genes. For example, *Per1, Per2 and Per3* are represented by a single *Per* gene. Among the corresponding proteins *PER, CRY and REV-ERB* proteins act as transcriptional repressors and ROR, CLOCK and BMAL1 as transcriptional activators. The central component of the model, the CLOCK_BMAL1 heterodimer, binds to the promoter regions of other clock genes (*Per, Cry, Ror, Rev-erb*) activating their transcription. We posit a reversible phosphorylation mechanism for all cytosolic circadian clock proteins in generating the model, as phosphorylation plays a regulatory role in the circadian system. The reversibility of the phosphorylation allows for dynamic changes in protein activity and regulation of circadian processes. This hypothesis is based on evidence that phosphorylation of key clock proteins (e.g. PER, CRY, BMAL1) alters their stability, localization, protein-protein interactions, and transcriptional activity over the circadian cycle^28,29^. The ROR_c_ and REV-ERB_c_ proteins (where the subscript ‘c’ stands for ‘cytosolic’) are phosphorylated reversibly, following which they are transported to the nucleus. In the nucleus, ROR_n_ and REV-ERB_n_ (the subscript ‘n’ stands for ‘nuclear’) bind to the promoter region of Bmal1 to regulate its transcription with ROR_n_ serving as an activator and REV-ERB_n_ a repressor. Likewise, *Bmal1* is translated to BMAL1_c_ protein in the cytosol and thereafter phosphorylated reversibly. BMAL1_c_ is transported to the nucleus where it forms a reversible heterodimer with the CLOCK protein to generate a protein complex, we term “CLOCK_BMAL1”. The PER and CRY proteins in the cytosol are phosphorylated reversibly, upon which the unphosphorylated PER and CRY proteins form the reversible PER-CRY_c_ heterodimer which is then transported into the nucleus. PER-CRY_n_ represses the transcription of the genes *Per* and *Cry*, as well as *Reverb* and *Ror*, by binding to CLOCK_BMAL1, thus suppressing the activity of CLOCK_BMAL1. The dissociation of the transcription factor PER-CRY from the promoters of these genes allows CLOCK_BMAL1 to activate transcription of *Per, Cry, Ror, Rev-erb,* thus starting the process all over again. We summarize these interactions with a wiring diagram that consists of two main (positive and negative) coupled feedback loops (Fig 2B).

**Figure 2:**
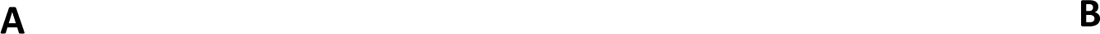
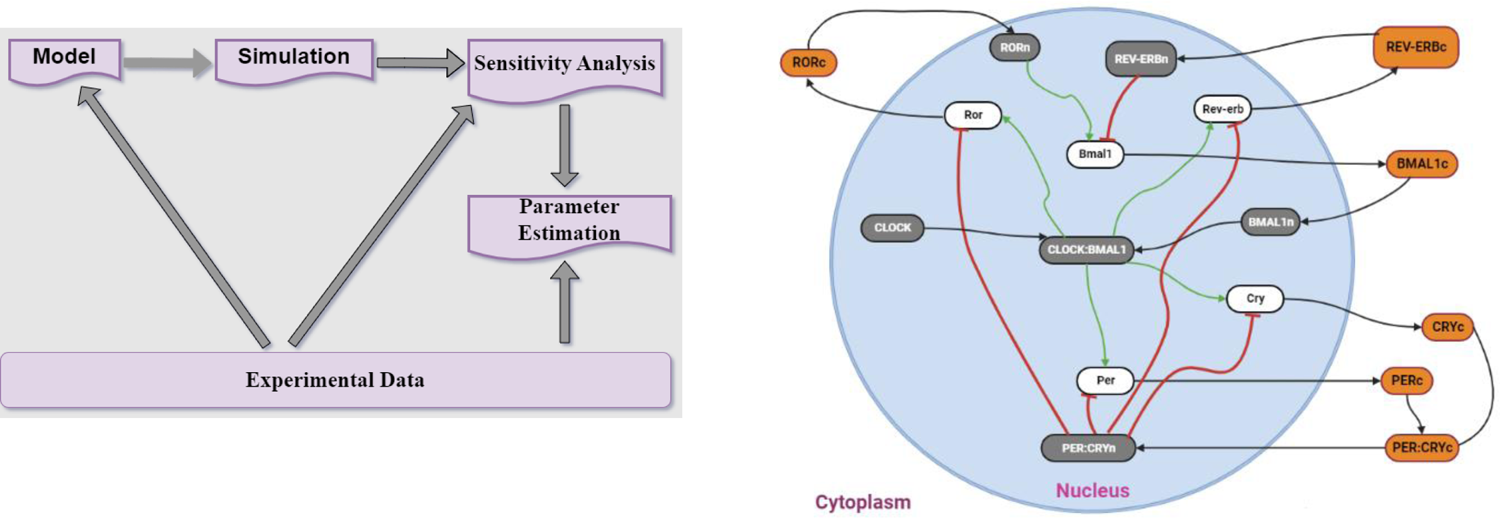
Modeling framework and schematic network diagram. **(A)** Construction of a deterministic model of the mammalian circadian clock, and data-driven sensitivity analysis and parameter estimation. **(B)** Schematic diagram of the principal positive and negative regulatory feedback loops in the mammalian clock network. The core negative feedback loop is formed by PER:CRY heterodimers that repress their own transcription. Another regulatory (positive and negative feedback) loop is formed by *Ror and Rev-erb* competing for binding to ROR/REV-ERB-response element (RORE) to regulate *Bmal1*. Green lines indicate activation by the transcription factors CLOCK-BMAL1 and RORn, red lines indicate repression of genes by transcription factors, and black lines indicate the translation and translocation of mRNAs and proteins. White ovals represent genes, gray ovals proteins in the nucleus and orange ovals proteins in the cytosol.

### Multicellular spatiotemporal model of the core clock genes exhibits coupling among cells

Gene regulatory networks can be described by a set of coupled ordinary differential equations (ODEs), where molecular interactions are described by Michaelis-Menten or higher order Hill functions^30^. The regulatory network in Fig 2B was translated into two deterministic spatiotemporal models (Supplementary Material:Method): one without intercellular communication (coupling), and one with coupling. We used Michaelis-Menten and Hill function approximations for all transcription factor – gene interactions, and assumed mass action and linear kinetics for degradation, translation, and complex formation. In absence of synchronization signals, autonomous cells in the suprachiasmatic nucleus or other peripheral tissues like the liver would oscillate with different periods, amplitudes, and phases. However, intercellular communication induced by extrinsic or intrinsic signals ensures synchrony in oscillations and resulting biological functions^21,27,31^ The two models were developed to investigate the spatiotemporal expression of circadian clock genes in the liver. We used a dataset^18^consisting of single-cell RNA seq and single molecule fluorescence in situ hybridization (smFISH), which found that expression of the core circadian clock genes in the liver lobule was non-zonated due to coupling among autonomous cell oscillators^18^. For the uncoupled model, the wiring diagram in Fig 2B was translated into a system of 23 differential equations describing gene transcription and translation, protein complex formation, phosphorylation, and inhibition. We applied this model to an assembly of over 435 cells laid out spatially in the form of a liver lobule (Lobule geometry described in a piff file in Compucel3d simulations). We assumed a Gaussian distribution to randomize the transcription rate parameters for every variable in each cell. This stochastic modeling approach aimed to capture potential heterogeneity across cells in transcription rates arising from the desynchronization in gene expression. The spatial and temporal expression for each gene was then simulated in the Compucell3d^32,33^ modeling environment.

The simulated mean expression of both *Per* and *Bmal1* genes at timepoints 6,12,18, and 24 hrs. was not zonated across the portal-central axis (Fig 3A), confirming results from previous studies^18^. The temporal profile for both genes in the uncoupled model showed autonomous oscillations across the lobule with period ranging from 21 to 28 hours and amplitude from 2.8 to 4.7 for *Per* and period ranging from 21 to 28 hours and amplitude from 0.8 to 3.2 for *Bmal1* (Fig 3B, 3E and 3F). For the coupled model, we assumed synchronization of autonomous cells across the liver lobule is achieved by communication (mediated by a putative coupling ligand) between each cell and its neighbors. The transmission of the coupling ligand across the liver lobule effectively synchronizes the period of the oscillation^31^. We assume that global synchronization across the lobule is achieved through the average concentration of the coupling ligand affecting clock genes via receptor molecules^21^. Data from Finger et al^21,27^ revealed a potential coupling factor; the transforming growth factor-beta (TGF-β), activates early transcription factors to control the molecular clock machinery, leading to a significant up-regulation of *Per 2* mRNA levels after 2-4 hrs. The spatial and temporal expression for each gene was then simulated in Compucell3d, showing coupling across the portal-central axis (Fig 3C). The temporal limit cycle oscillation profile for both genes showed coupled oscillations with period 24-25 hours and amplitude of 5.2 for *Per* and period 23-25 hours and amplitude 2.2 for *Bmal1* (Fig 3D, 3E and 3F).

**Figure 3.**
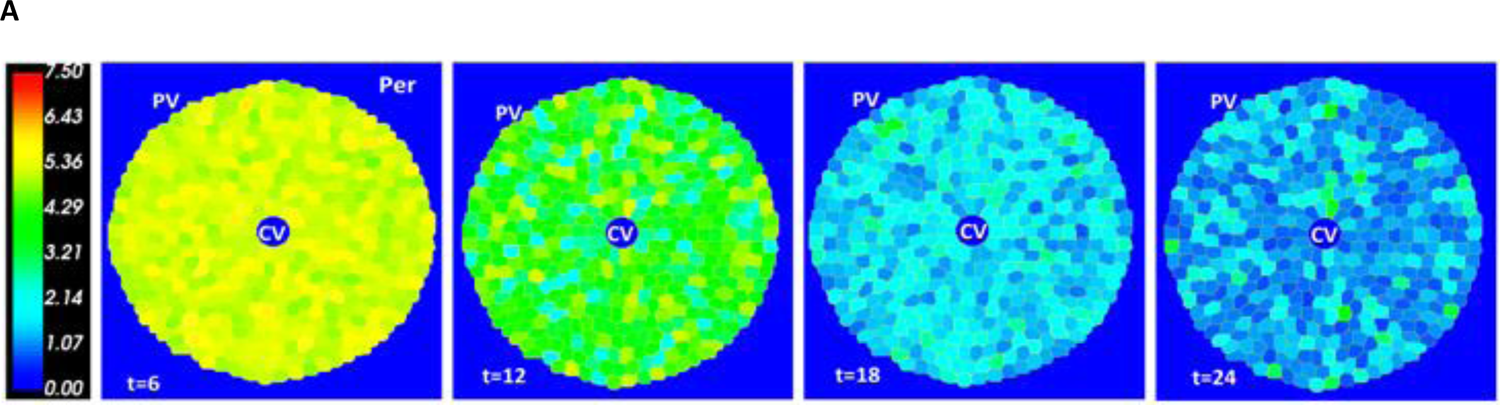
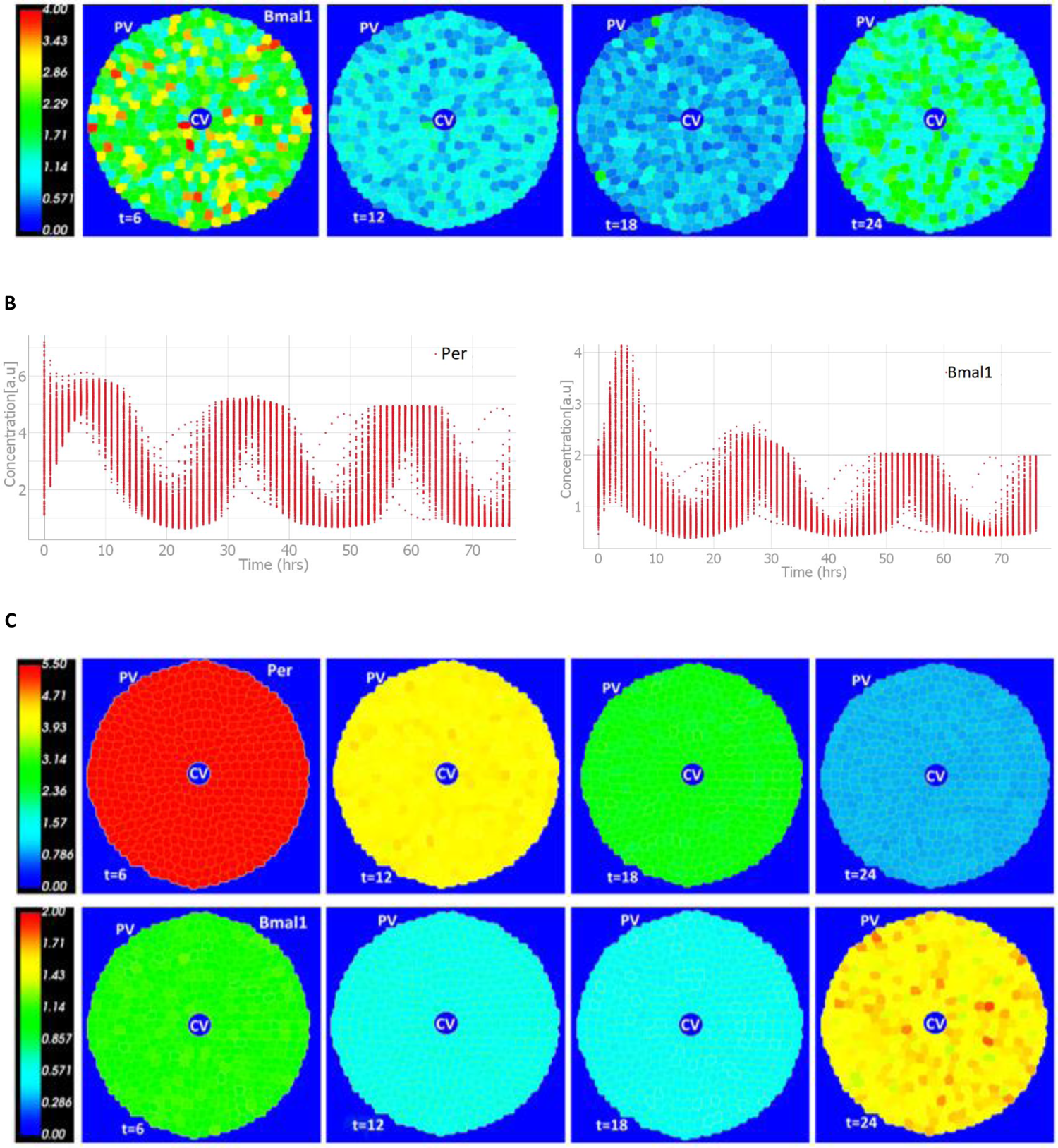
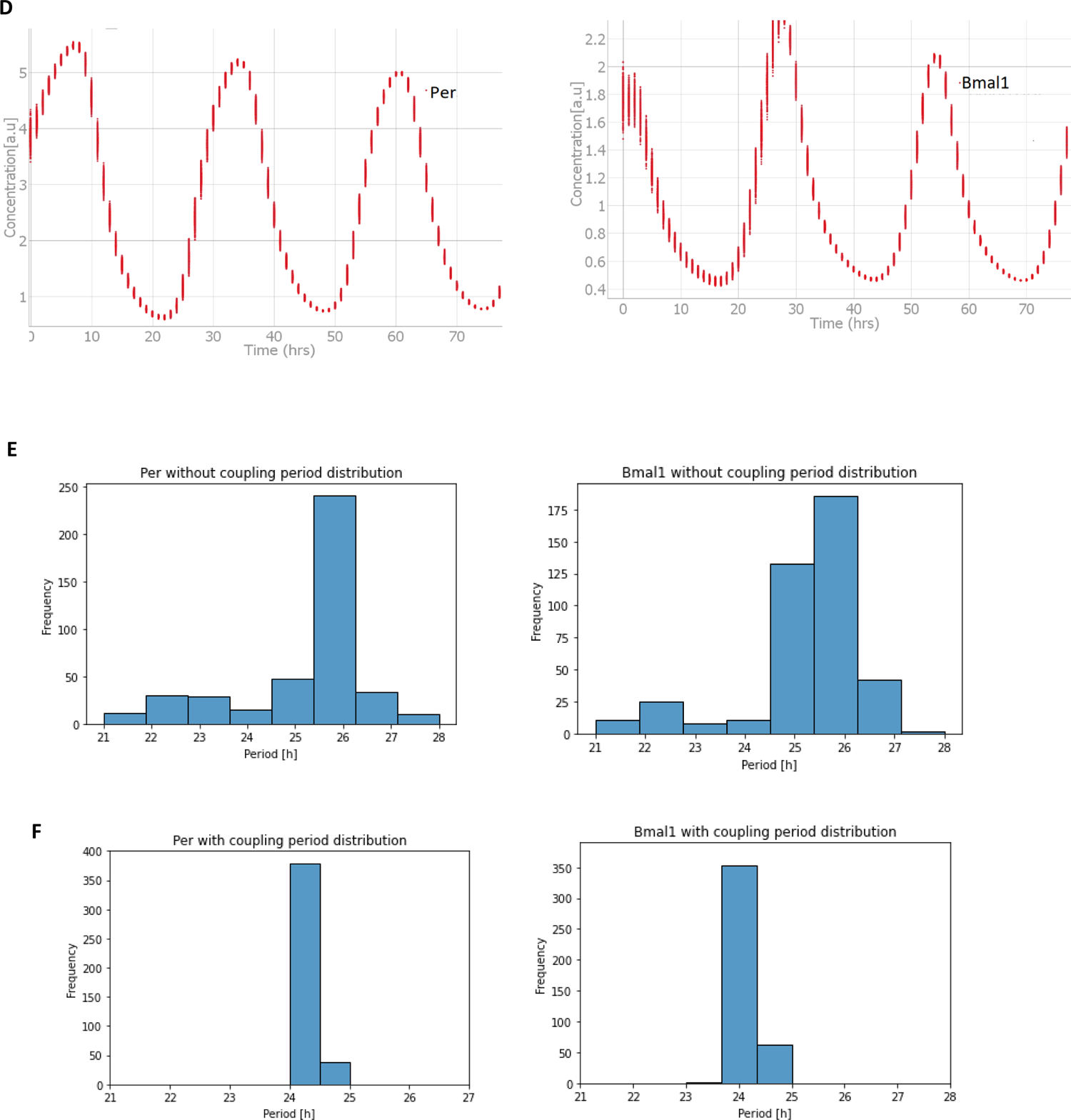
Multicellular spatiotemporal model simulations. Expression across the liver lobule of **(A)** Period gene (*Per*) and Brain and Muscle ARNT-Like 1 (*Bmal1*) without coupling at circadian times ZT 6, 12, 18, and 24 hours. The periportal lobule displays non-zonated mean expression. as reported in Droin et al^18^. **(B)** Limit cycle oscillations of the *Per* and *Bmal1* genes for 435 cells without coupling with varying period and amplitude values for *Per* (period: 21-28 hours, amplitude: 2.8-4.7) and *Bmal1* (period: 21-28 hours, amplitude: 0.8-3.2). **(C)** The spatial expression of *Per*, and *Bmal1* genes for 435 cells with coupling at circadian time ZT 6 12, 18 and 24 hours across the liver lobule. The expression profile shows a synchrony across all time points. **(D)** Limit cycle oscillations of the *Per* and *Bmal1* genes with coupling producing synchronized period and amplitude for *Per* (period: 24-25 hours, amplitude: 5.0-5.2) and *Bmal1* (period: 23-25 hours, amplitude: 2.0-2.2). The color bar shows the expression values of genes in each cell across the central (CV) and portal (PV) in the liver lobule. Each dot represents the time-dependent expression of a gene per cell. **Distribution of uncoupled and coupled individual cell periods. (E)** The distribution of uncoupled Per and Bmal1 oscillation periods for over 400 cells in the model lobule. The period ranges from 21 hours to 28 hours. **(F)** The distribution of coupled Per and Bmal1 oscillation periods for over 400 cells in the model lobule. The period ranges between 24 hours to 25 hours for Per oscillations and 23 hours to 25 hours for Bmal1 oscillations.

### Sensitivity Analysis

Next, we carried out parameter sensitivity analysis on the model dynamics (period of the oscillation, the amplitude of the oscillation, the phase and the average amount of mRNA produced) using the AMIGO2 toolbox ^34,35^ in MATLAB. Specifically, we tested the effect of the transcription and degradation rate parameters on the model dynamics. We adopted the Latin hypercube sampling method (LHS) which is noted to yield more precise estimate to select the number of intervals to divide the tested range of the variable into.

### The transcription and degradation rate parameters of the Reverb gene respectively show negative and positive correlation with the average expression level of *Bmal1*

We investigated the effect of the transcription and degradation rate parameters on average expression of the observables (mRNA levels of clock genes) for a complete circadian cycle (24 hrs) after reaching a stable oscillation. A 10% increase in the transcription rate of each gene demonstrated a direct correlation with its corresponding mRNA level, as raising the transcription rate parameter resulted in a rise in the average expression level of the associated observables. (Fig 4B). The positive and negative feedback regulations in the gene network were also reflected in the sensitivity analysis of the transcription and degradation rate. An increase in the transcription rate parameter of the Cry gene, vCs, decreased the average expression of Per mRNA because of the delay inhibition by PER-CRYn. Likewise, an increase in the transcription rate parameter of Bmal1, vBs, deceased the average expression of Reverb and increased the average expression of Per and Cry because of the positive feedback loop between Bmal1, Per, and Cry, and the negative feedback loop between Bmal1 and Reverb (Fig 4A-4B). Conversely, a 10% increase in the degradation rate parameters exhibited an inverse correlation with their corresponding observables, resulting in a decline in the average expression level of their associated genes. (Fig 4C). Also, an increase in the degradation rate parameter of Bmal1, d_mB, led to an increase in the expression level of the Reverb gene due to the negative feedback loop between Bmal1, PER-CRY and Reverb genes. Other relations between the degradation rate parameters and observables are shown in Fig 4C. The clock gene network demonstrated greater sensitivity to transcription rate versus degradation rate changes. Of all genes analyzed, only Bmal1 was sensitive to degradation rates of the entire network, despite being directly regulated only by Ror and Reverb.

**Figure 4:**
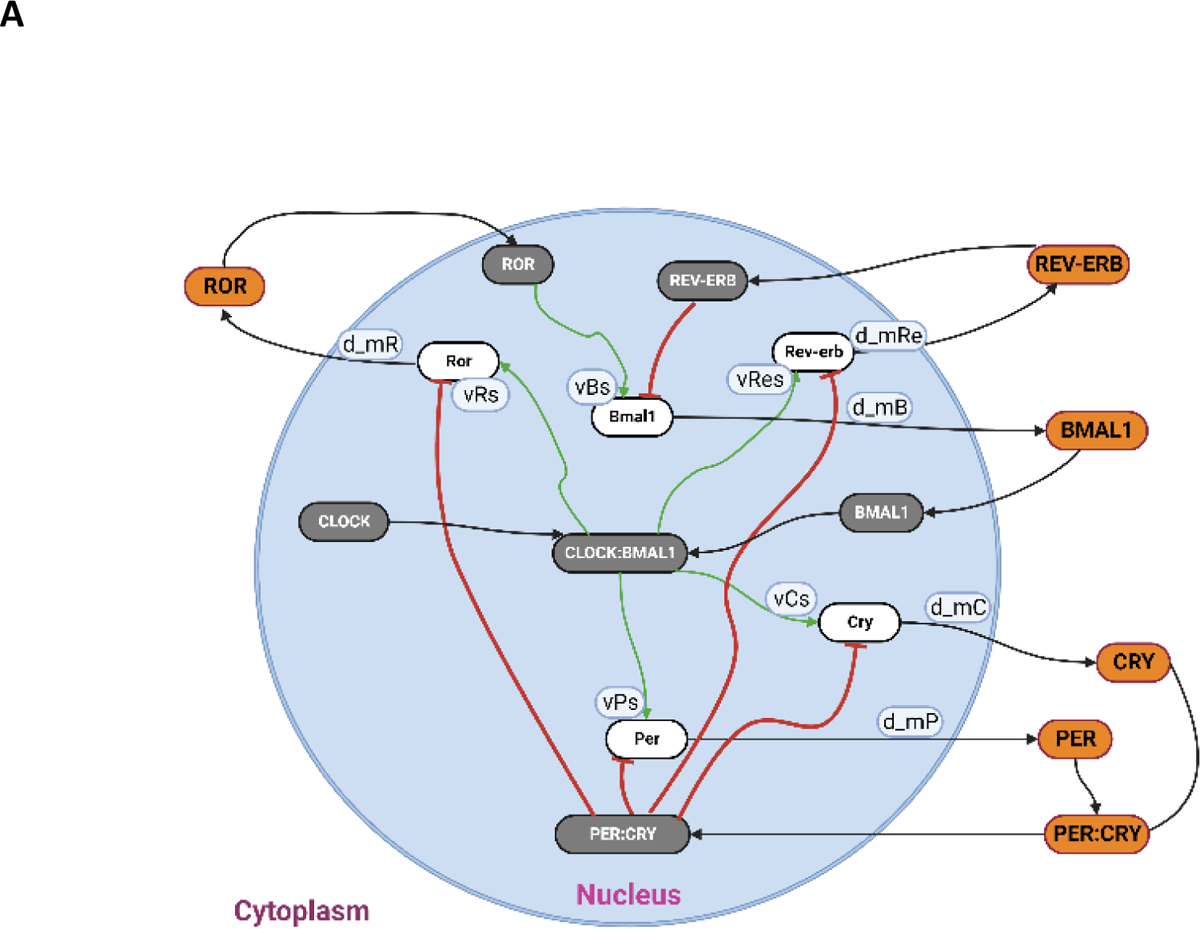
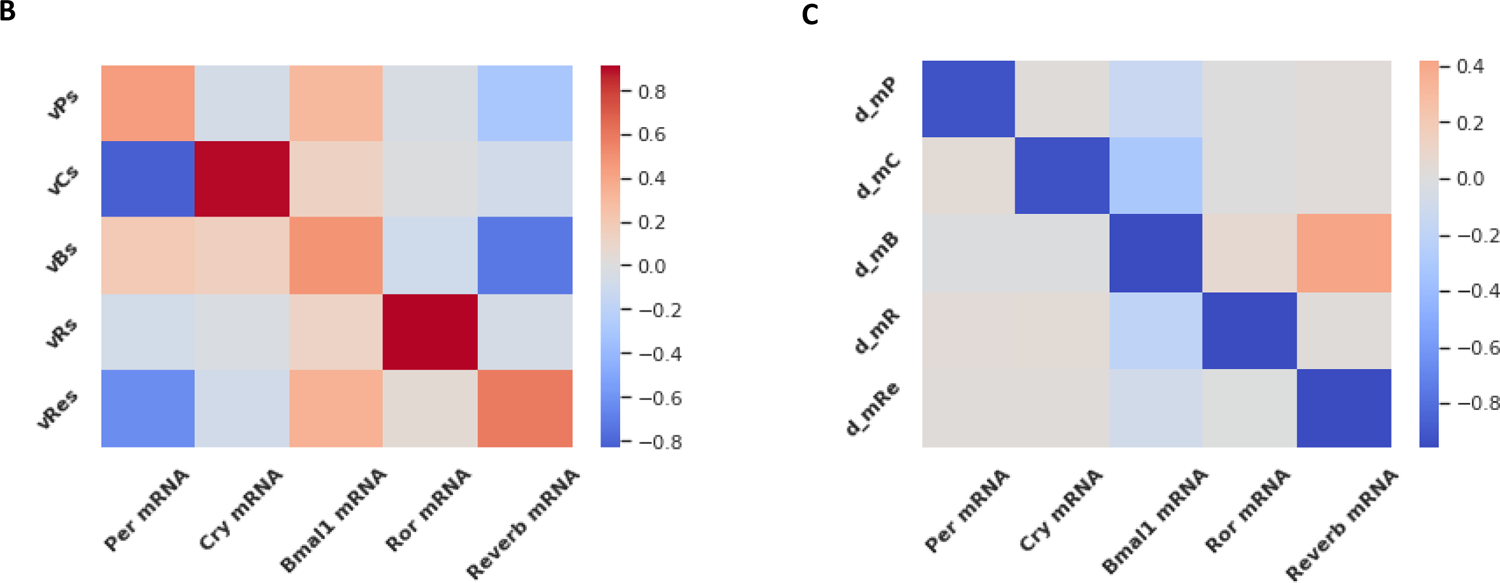
(A) Schematic diagram of the mammalian clock network with its transcription and degradation parameters. (B) Sensitivity analysis of transcription rate parameters. The transcription rate parameter for the observables (mRNA levels of clock genes) were optimized and the dynamics revealed a positive correlation between the transcription rate parameters and the average mRNA level of the respective gene. **(C) Sensitivity analysis of degradation rate parameters.** Optimization of the degradation rate parameter for the observables (mRNA levels of model clock genes) revealed a negative correlation between the degradation rate parameters and their respective average mRNA. Also, an increase in the degradation rate parameter of the Bmal1 gene, d_mB, led to an increase in the expression level of the Reverb gene due to the negative feedback loop between Bmal1, PER-CRY and Reverb gene. Additional correlations between the degradation rate parameters and other observables were demonstrated.

### Increase in the transcription rate parameters lead to a non-monotonic decrease in the oscillatory period while increasing the amplitude of the oscillation

The observed oscillatory period is the same for all circadian clock genes, ranging from 23.5 hrs to 24.5 hrs ^12^. The amplitude which was measured as the absolute difference between the peak and the trough levels of the oscillation on the other hand varies across the genes due to different biochemical reactions involved in each gene transcription. A sweep on the transcription rate parameters within the range of 0-5 showed a non-monotonic change in the oscillatory period. The period of the circadian clock genes, *Per* and *Reverb,* decreased monotonically with an increase in the transcription rate parameter for *Per* and *Reverb*, vPs and vRes, after an initial linear increase at very low transcription rates (Fig 5A). We observed a monotonic increase in the oscillation period of the *Ror* gene with an increase in its corresponding transcription rate parameter (result not shown). This was in line with previous studies^36^ attributing this dependence to the positive feedback loop mediated by ROR. Using the same parameter values (0-5) we investigated the effect of the transcription rate parameters on the amplitude of the circadian clock oscillation. We observed a monotonic increase in amplitude across all clock genes in the model with respect to an increase in their corresponding transcription rate parameter (Fig 5B).

**Figure 5:**
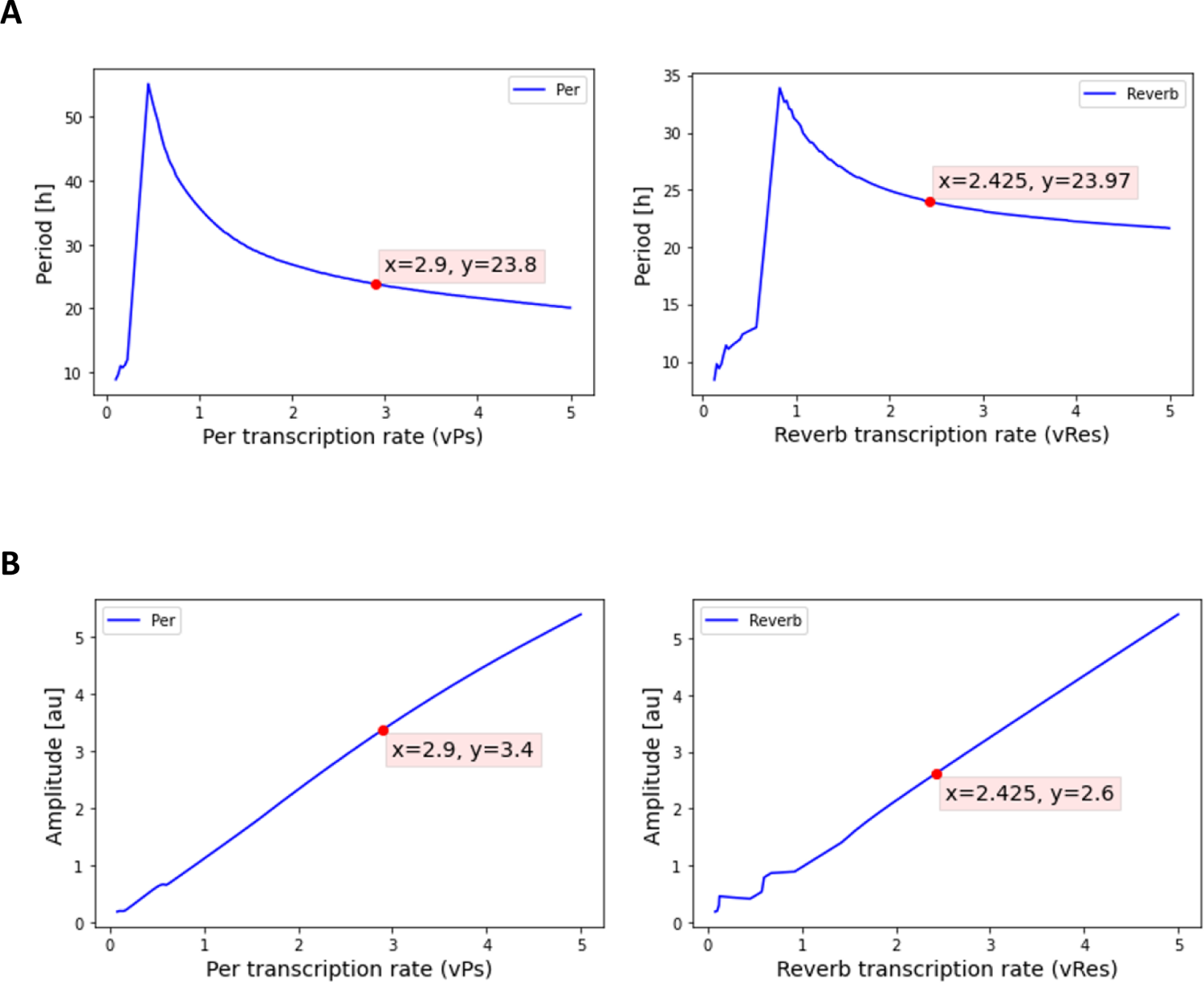
A non-monotonic period and amplitude dynamics for observables *Per* mRNA and *Reverb* mRNA as gradient of the transcription rate parameters applied in the model. (A) Oscillation period changes with respect to changes in transcription rate parameter for Per mRNA and Reverb mRNA. The oscillatory period exhibits a monotonic decrease with an increase in the transcription rate parameter. The red dot corresponds to the wild-type parameter value for the transcription rate parameter for Per and Reverb (x=2.9 and x=2.425) respectively, and their corresponding oscillatory period (y=23.8 h and y=23.97) (**B**) Oscillation amplitude changes with respect to changes in transcription rate parameter for Per mRNA and Reverb mRNA. A monotonic increase of oscillatory amplitude is observed with an increase in the transcription rate parameter. The red dot corresponds to the wild-type parameter value for the transcription rate parameter for Per and Reverb (x=2.9 and x=2.425) respectively, and their corresponding oscillatory amplitude (y=3.4 and y=2.6)

### The period undergoes a concave change (with a downward slope followed by an upward slope), and a monotonic decrease in amplitude as the degradation rate parameter increases for *Per* and *Reverb*

We conducted a parameter sweep on the degradation rate parameters to explore their impact _on both the period and amplitude (_Fig 6_)_. We observed a concave change in the oscillatory period of the circadian clock genes with respect to an increase in their respective degradation rate parameters (range 0 to 1.0). The period first decreases, then increases monotonically with an increase in the degradation rate parameters (Fig 6A). The period of the circadian clock genes, *Per* and *Reverb*, likewise showed concave period changes with an increase in the degradation rate parameter for Per and Reverb, d_mP and d_mRe respectively (Fig 6A). We observed a monotonic decrease in amplitude across all clock genes in the model with respect to an increase in their corresponding transcription rate parameter (Fig 6B).

**Figure 6:**
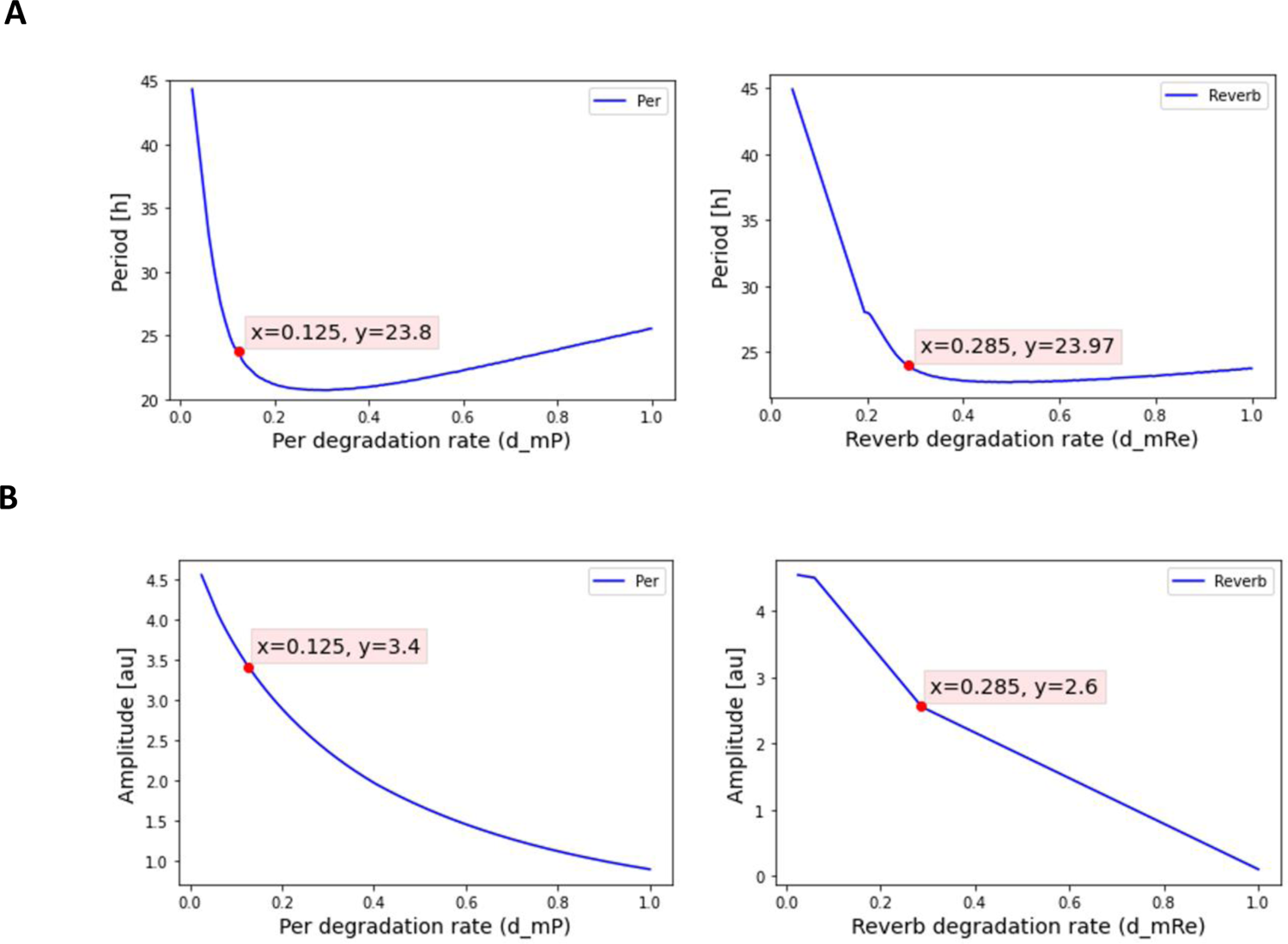
A concave dynamic period for *Per* and *Reverb* as degradation rate parameter increases. **(A)** The oscillation period of both *Per* and *Reverb* decreases and increases giving a concave dynamic with respect to an increasing degradation rate parameter. **(B)** Oscillation amplitude for *Per* and *Reverb* observed a monotonic decrease with an increase in the degradation rate parameter. The red dots in **(A)** and **(B)** denote their degradation rate parameter value and the resulting period/amplitude value respectively.

### Bifurcation analysis shows that *Cry* and *Ror* exhibit Hopf bifurcations with varying transcription rate parameter value

Next, we carried out bifurcation analysis of the dependence of oscillatory rhythm dynamics (period and amplitude of oscillation, the phase and the average amount of mRNA produced) on parameter values in XPPAUT^37^. The bifurcation diagram, based on one parameter, revealed a dependence of both amplitude and periodic solutions on the transcription rates (vCs for Cry mRNA and vRs for Ror mRNA). The bifurcation diagram generated display a cyclic oscillation, with stable and unstable limit cycles depicted in green and blue, respectively, indicating Hopf bifurcation (HB) points (Fig 7 A&C). Next, by extrapolating from the bifurcation diagram, we derived the stable and unstable period oscillation for both Cry and Ror (Fig 7 B&D). The alignment of the stable and unstable parameter values in both period and limit cycle oscillation for each gene was evident. Subsequently, we extended the bifurcation analysis to include degradation parameters of observables, demonstrating the stability of the model’s rhythmic dynamics (Supplementary figures).

**Figure 7:**
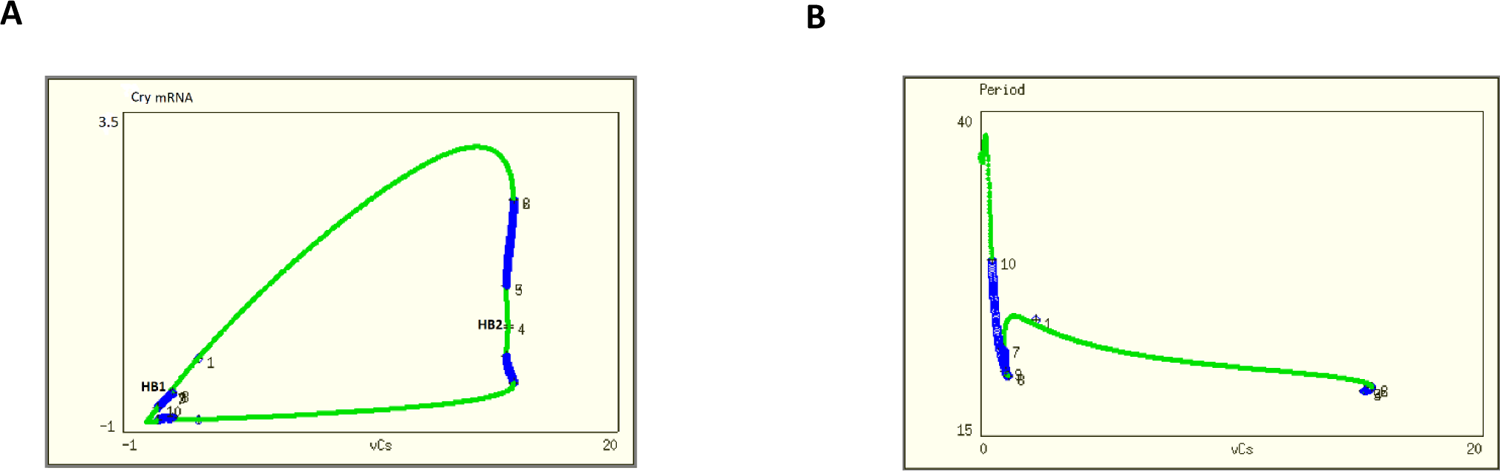
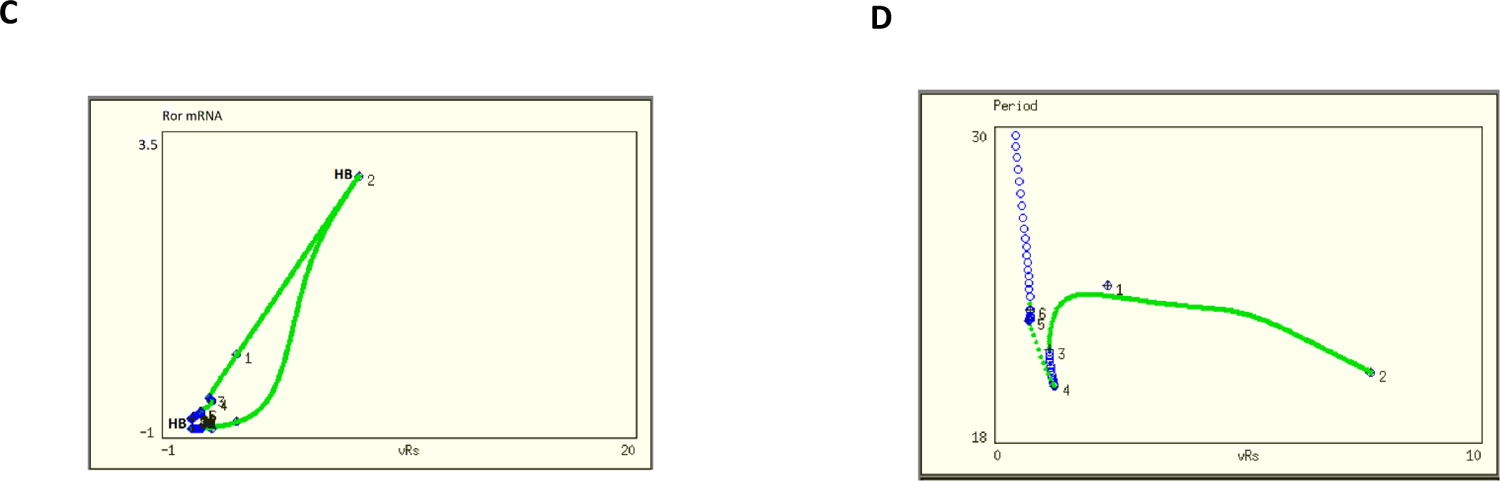
One parameter bifurcation analysis of the model rhythmic dynamics. **(A&C)** The bifurcation diagram of the transcription rate parameter of Cry mRNA, (vCs) and Ror mRNA, (vRs). The green and blue lines represent stable unstable limit cycles respectively. HB shows the Hopf bifurcation points in the bifurcation diagram. **(B&D)** Bifurcation diagram showing the periodic stability of the transcription rate parameter of Cry mRNA, (vCs) and Ror mRNA, (vRs). The green line represents stable period and the blue line represent unstable period of the model.

### Model Parameter Estimation

Using experimental data from hepatic single nuclei RNA-sequence data obtained from male C57BL/6 mice housed with a 12:12 light:dark cycle to synchronize with the environmental circadian cycle, we employed the parameter estimation algorithm in AMIGO2^34,35^. The algorithm was used to optimize transcription and degradation rate parameters, aiming to fit the model prediction to the hepatic single nuclei RNA-sequence experimental data. The distance between the model predictions and experimental data, as quantified by the cost function, depends on how well the parameter estimation algorithm optimizes agreement between the data and predictions. Specifically, the algorithm aims to minimize the cost function, which measures the discrepancy between the experimentally observed values and the values predicted by the model using a given parameter set. The smaller the overall distance as calculated by the cost function, the better the parameter set matches the available data. We defined our cost function as a maximum (log-) likelihood function due to the availability and the nature of the measured noise from the experimental data. The initial conditions for the transcription and degradation rate parameters that were estimated started at the wild-type values used in the model. Upper and lower bounds for these parameters were set based on values reported in the literature on circadian system modeling ^38–40^. We mathematically define a non-linear programming problem (NLP) with algebraic constraints to find transcription and degradation rate parameters that minimize the cost function.

### Estimation of transcription and degradation rate parameters

To evaluate our model’s prediction accuracy, we used single-nuclei gene expression data from male C57BL/6 mice at 2, 4, 8, 12, 18 and 24-hour timepoints. We considered the core circadian genes used in our model: *Per, Cry, Bmal1, Ror*, and *Reverb*. Normalized RNA-seq data replicates for these genes represented the experimental dataset for optimizing our model’s transcription and degradation rate parameters. We averaged isoform expression to represent overall gene expression. We averaged expression levels of gene isoforms to represent each gene. Given our large non-linear model, local optimization risks converging to local minima. We used the enhanced Scatter search eSS hybrid global-local optimization algorithm in the AMIGO toolbox to efficiently find an accurate global optimum solution^41,42^. We optimized five transcription and five degradation rate parameters, each tied to an observable in our model. The optimized parameters positively correlated with the nominal wild-type parameters. Consequently, the model predictions using the optimized parameters positively correlated with the experimental measurements for all five observables (Fig 8 A&B). Optimizing the transcription and degradation rates for each observable improved the parameter estimates, while maintaining the rhythmic patterns of oscillation phase, acro-phase, amplitude, and period (Fig 8 C-F). The results show an improvement in the parameter estimates. The model results maintained the rhythmic pattern from the experimental data which include the phase of the oscillation, the acro-phase of the oscillation (timepoint with maximum expression), the amplitude and period of the oscillation. However, due to the lack of protein data, particularly in the cytoplasm and nucleus, perfect optimization of the model was not achieved, suggesting potential enhancement with complete protein data availability.

**Figure 8:**
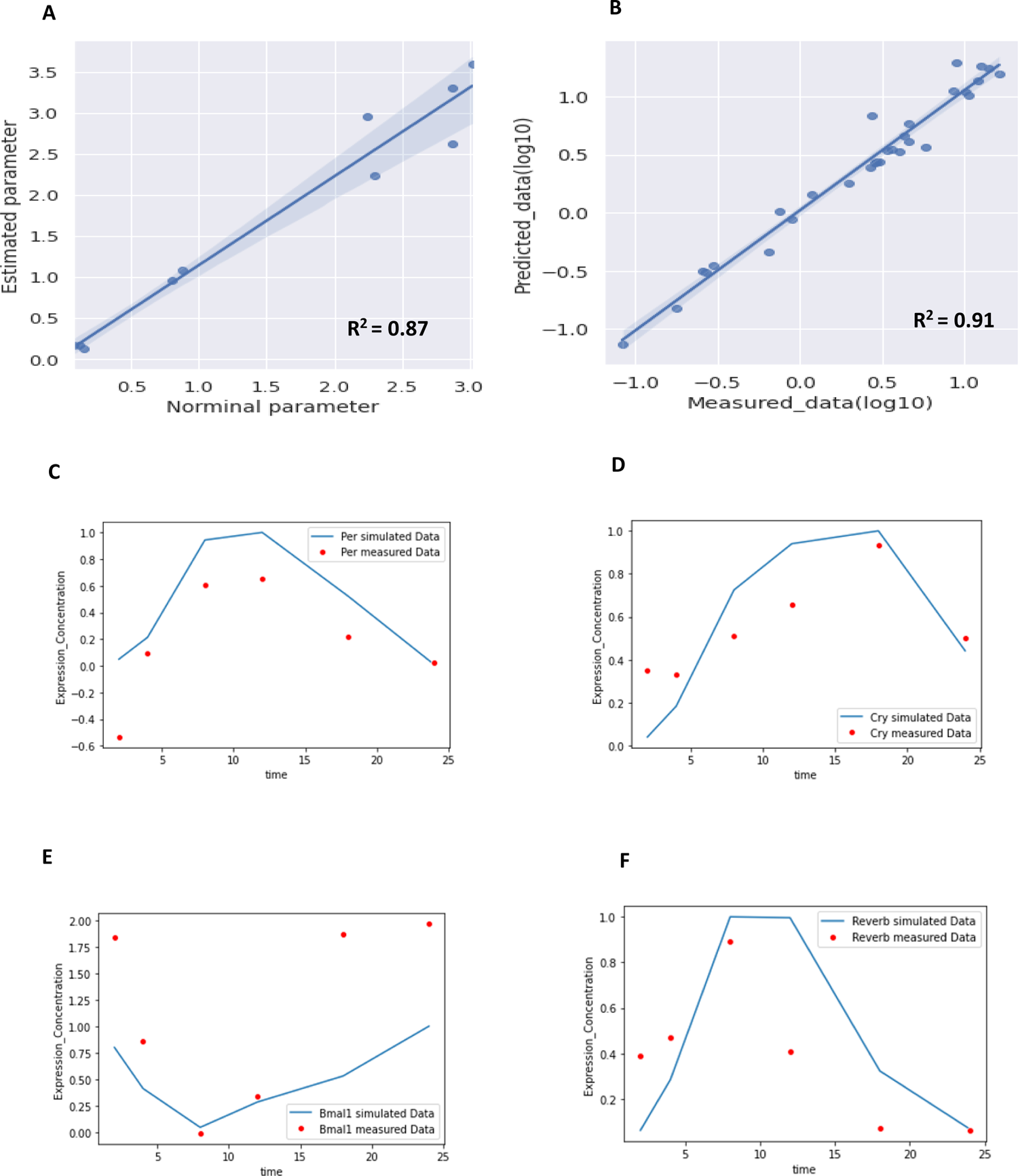
The predicted model results and experimental data fit for observables using maximum (log-) likelihood function. (A) Correlation between predicted transcription and degradation and wild type transcription and degradation rate parameter values. **(B)** Correlation between model predictions and measured data. **(C-F) The fitted curves for the model variables (**Per, Cry, Bmal1 and Reverb). The red dotted line indicates the experimental data at timepoints 2,4,8,12,18 and 24 h. The solid blue line indicates the predicted results from using maximum (log-) likelihood function to fit the model to the experimental dataset.

## Discussion

Circadian rhythms display an endogenous, entrainable oscillation with a period of about 24 hours, allowing organisms to anticipate and adapt to daily environmental changes in light, temperature, and nutrient availability^43^. Dynamic modeling of these biological processes from the gene regulatory to multicellular network levels provides key insights into the fundamental properties, physiology, and behaviors related to circadian rhythmicity^44^. Though experimental and theoretical explorations have extensively detailed the circadian clock gene regulatory network^45^, few studies have examined the cell-cell communication processes enabling synchronization of circadian period, amplitude, and phase between autonomous cellular oscillators. Elucidating circadian synchronization through cell-cell coupling can advance our understanding of circadian rhythm robustness and plasticity at tissue level^18^. Neurotransmitters, acting as coupling factors, have been shown to regulate the synchronization mechanism in the suprachiasmatic nucleus (SCN)^31,46^. However, the coupling factors in other peripheral tissues are unknown. In this study, we developed mathematical models of the mouse hepatic circadian clock to examine intercellular communication and synchronization of autonomous oscillators across the murine liver lobule. The models incorporated core clock genes and their regulatory interactions in individual hepatocytes. Simulations showed that incorporating cell-cell coupling led to synchronized gene expression between hepatocytes, matching experimental findings^18,47^. Strong synchronicity of circadian oscillation has been associated with period lengthening. However, our models suggest that without synchronization the period is variable outside of the near 24-hr range. Therefore, optimal cell-cell coupling is required to achieve both synchronicity between cells and appropriate oscillatory periods (Fig 3E and 3F).^31,48^. Synchronicity of circadian oscillation has been shown to induce key cell-cycle events including cyclin-dependent kinase network activation, cell growth, DNA replication, and cytokinesis^49^. A weak synchronicity at the cellular level in the Sensitivity analysis revealed dependencies between clock components; for instance, increased Per transcription decreased Cry expression, likely due to their mutual repression^51^. Specifically, increased *Cry* transcription rates markedly diminished *Per* mRNA levels. However, perturbations in the Per transcription rate did not comparably suppress Cryptochrome transcripts^40^. This aligns with experimental reporter assays demonstrating the repression strength of PER proteins on Clock/Bmal1-driven transcription is weaker relative to CRY^52^. The disproportionate parametric sensitivities suggest CRY dynamics play a more dominant role than PER in governing circadian rhythmicity via interlocking negative feedback loops. Validating the coupled mechanism in the model, we used bifurcation diagrams to show the periodic stability of the transcription rate parameters. Our theoretical bifurcation diagrams generate experimentally testable hypotheses, potentially through overexpression or knock down of circadian factors in vivo and in vitro to validate predicted changes in periodicity, amplitude, or phase shifts^36,53^.

Our knowledge of the complex mammalian circadian clock mechanism is still incomplete. More than 40 genes directly interact with the core clock genes in generating the circadian oscillatory rhythm^54^. It is therefore essential to understand the temporal and spatial dynamics and the regulatory mechanism of the circadian clock oscillation. We address this gap in knowledge with a detailed mathematical model incorporating known clock and associated rhythmic genes. The current model focused on healthy cells under normal circadian entrainment. An important extension would be to model circadian disruption by genetic alterations or toxicant exposure. The model could be used to predict effects of parameter changes representing mutations or cellular damage. Linking the circadian clock model with zonated metabolism models could offer insights into compounding hepatic effects across scales.

In conclusion, we have introduced two sets of mathematical models of the mouse hepatic clock, with and without the synchronization of cells in the hepatic lobule. We found that like the coupling of autonomous circadian oscillators in the suprachiasmatic nucleus (SCN), hepatic clock rhythms are also likely synchronized by a yet unknown coupling factor. Sensitivity analysis, bifurcation analysis, and parameter estimation from our model have shed more understanding on the physiology of the hepatic clock and ways it can be altered. The model demonstrated how variations in impact parameters influence the positive and negative feedback loops. Overall, the modeling framework presented here establishes the foundation for investigating circadian regulation and dysfunction in liver physiology and disease states.

## Author Contributions

D.M., and S.B. designed the study. D.M., D.F., and O.K., J.P.S., and Z.Q. developed the mathematical models and generated all the figures. D.M., S.B., O.K., D.F., Z.Q., and S.L., J.P.S. wrote the manuscript. All authors reviewed the manuscript.

## Data availability

The Single-nuclei RNA seq dataset used for this study is available from the corresponding Authors, D.M or SB, upon reasonable request. The codes used for generating the liver lobule geometry and all analysis are available at https://github.com/BhattacharyaLab

## Acknowledgements

This work was supported in part by NIH grants P42ES004911, the Superfund Research Program NIEHS SRP P42ES004911, R01ES031937, and Michigan State University AgBioResearch. JPS was supported by NIH grant U24 EB028887.

