## Supplementary material for "A multiscale model of the mammalian liver circadian clock supports synchronization of autonomous oscillations by intercellular communication": Method

### METHODS

#### MODEL DESIGN

This model was developed to study the communication between single cellular circadian oscillations in the liver lobule. The communication which is termed as 'coupling' in other studies synchronizes the several autonomous circadian oscillations to produce a coherent output in the suprachiasmatic nuclei (SCN) and other peripheral tissues. We base this model on cellular oscillators found in the liver lobule. Identifying and assembling relevant information, key interactions and connections between the clock gene network was achieved from literature. Five protein known clock genes were used in developing this model. For simplicity, we do not distinguish between the mutations in theses genes. Example *Per1*, *Per2* and *Per3* are represented by a single *Per* gene. The same principle applies to the proteins and respective protein complexes. Gene network used in developing this model is (*Per*, *Cry*, *Ror*, *Rev-erb* and *Bmal1*). This model emphasis on the grouping of the gene network (Activators and Repressors). The *Per*, *Cry*, and *Rev-erb* genes as the repressors and the *Ror*, and *Bmal1* genes as the activators. The interactions and connections between these gene network was then transformed to a wiring diagram that consist of two main dependent feedback loops: the *Per-Cry* negative feedback loop and the *Ror*, *Rev-erb*, *Bmal1* loop.

The central component of the model, *CLOCK_BMAL* transcription factor binds to the promoter regions of the clock genes (*Per*, *Cry*, *Ror*, *Rev-erb*) activating their transcription . These various *mRNA's* are translated into the respective proteins in the cytosol. The $ROR_c$ and REV-$ERB_c$ protein in the cytosol are phosphorylated reversibly. The unphosphorylated $ROR_c$ and REV-$ERB_c$ proteins are transported in the nucleus. In the nucleus, $ROR_n$ binds to the promoter region of the clock gene *Bmal1* activating its transcription. The *Bmal1 mRNA* is translated to $BMAL1_c$ protein in the cytosol which is phosphorylated reversibly. The unphosphorylated $BMAL1_c$ protein are transported in the nucleus where it forms a reversible heterodimer with the CLOCK protein to get the *CLOCK_BMAL* protein. The nuclear REV-$ERB_n$ protein binds to the promoter region of the *Bmal1* gene to repress the activity of the nuclear $ROR_n$ protein.The PER and CRY proteins in the cytosol are phosphorylated reversibly. The unphosphorylated PER and CRY protein in the cytosol form the reversible PER-CRY heterodimer which is then transported into the nucleus. The PER-CRY represses the transcription of their own genes (*Per* and *Cry* ) by binding with the *CLOCK_BMAL* protein. This association represses the transcriptional activities of all the genes that are activated by *CLOCK_BMAL*. The dissociation of the transcription factor *CLOCK_BMAL* from the PER-CRY protein allows the *CLOCK_BMAL* to bind to the promoter regions of the clock genes (*Per*, *Cry*, *Ror*, *Rev-erb*) and starting the process all over.

Two set of models were developed in this studies.(1) Model without communication between the cellular oscillators leading to a non-synchronized expression across the central to portal axis of the liver lobule and (2) model with communication between cells leading to synchronized expression across the central to portal axis of the liver lobule.

##### Model 1: Model without communication between the cellular oscillators

• The first five ordinary differential equation of this model represent the transcription of *Per*, *Cry*, *Bmal1*, *Ror* & *Rev-erb* genes into their respective mRNA's denoted by $M_P$, $M_C$, $M_B$, $M_{Ro}$ & $M_{Re}$ respectively by the nuclear *CLOCK_BMAL* protein ($CB_n$), *PER_CRY* protein ($PC_n$), *ROR* protein ($Ro_n$) and *REV_ERB* protein ($Re_n$).

$$\frac{dM_P}{dt} = light * \frac{V_{sp} * (CLOCK\_BMAL1)^n}{Kip^n + (CLOCK\_BMAL1 * PER\_CRY)^m + (CLOCK\_BMAL1)^n} - d_p * M_P \quad (1)$$

$$\frac{dM_C}{dt} = light * \frac{light * V_{sc} * (CLOCK\_BMAL1)^o}{Kic^o + (CLOCK\_BMAL1 * PER\_CRY)^p + (CLOCK\_BMAL1)^o} - d_c * M_C \quad (2)$$

$$\frac{dM_B}{dt} = \frac{V_{sb} * (ROR)^p}{Kib^p + (ROR * REV\_ERB)^q + (ROR)^p} - d_b * M_B \quad (3)$$

$$\frac{dM_R}{dt} = \frac{V_{sr} * (CLOCK\_BMAL1)^r}{Kir^r + (CLOCK\_BMAL1 * PER\_CRY)^s + (CLOCK\_BMAL1)^r} - d_r * M_R \quad (4)$$

$$\frac{dM_{Re}}{dt} = \frac{V_{sre} * (CLOCK\_BMAL1)^t}{Kire^t + (CLOCK\_BMAL1 * PER\_CRY)^u + (CLOCK\_BMAL1)^t} - d_{re} * M_{Re} \quad (5)$$

• The next six ordinary differential equations of this model represent the translation and reversible activities of the unphophorylated PER, CRY, BMAL1, REV-ERB, ROR protein and PER-CRY protein complex in the cytoplasm denoted by $P_c,\ C_c,\ B_c,\ Re_c$ , $Ro_c$ & $PC_c$ respectively.

$$\frac{dP_c}{dt} = k * Per\_mRNA + Kpc1{*}(CYTOSOLIC\_PER\_CRY\_PROTEIN) - Kpco{*}(CYTOSOLIC\_PER\_PROTEIN) * (CYTOSOLIC\_CRY\_PROTEIN) - Kpc * ((CYTOSOLIC\_PER\_PROTEIN)) + Kppc * (PHOS\_CYTOSOLIC\_PER\_PROTEIN) - dpc * (CYTOSOLIC\_PER\_PROTEIN) \quad (6)$$

$$\frac{dC_c}{dt} = k1 * Cry\_mRNA + Kpc1{*}(CYTOSOLIC\_PER\_CRY\_PROTEIN) - Kpco{*}(CYTOSOLIC\_PER\_PROTEIN) * (CYTOSOLIC\_CRY\_PROTEIN) - Kcc * ((CYTOSOLIC\_CRY\_PROTEIN)) + Kcpc * (PHOS\_CYTOSOLIC\_CRY\_PROTEIN) - dcc * (CYTOSOLIC\_CRY\_PROTEIN) \quad (7)$$

$$\frac{dB_c}{dt} = k2 * Bmal1\_mRNA - Kbcc{*}(CYTOSOLIC\_BMAL1\_PROTEIN) - Kbc{*}((CYTOSOLIC\_BMAL1\_PROTEIN)) + Kbpc * (PHOS\_CYTOSOLIC\_BMAL1\_PROTEIN) - dbc * (CYTOSOLIC\_BMAL1\_PROTEIN) \quad (8)$$

$$\frac{dRo_c}{dt} = k3 * Ror\_mRNA - Krcc{*}(CYTOSOLIC\_ROR\_PROTEIN) - Krc{*}((CYTOSOLIC\_ROR\_PROTEIN)) + Krpc * (PHOS\_CYTOSOLIC\_ROR\_PROTEIN) - drc * (CYTOSOLIC\_ROR\_PROTEIN) \quad (9)$$

$$\frac{dRe_c}{dt} = k4 * Rev\ _erb\ _mRNA - Krecc * (CYTOSOLIC\_REV\_ERB\_PROTEIN) - Krec * ((CYTOSOLIC\_REV\_ERB\_PROTEIN)) + Krepc * (PHOS\_CYTOSOLIC\_REV\_ERB\_PROTEIN) - drec * (CYTOSOLIC\_REV\_ERB\_PROTEIN) \quad (10)$$

$$\frac{dPC_c}{dt} = Kpco * ((CYTOSOLIC\_PER\_PROTEIN) * (CYTOSOLIC\_CRY\_PROTEIN)) - Kpcc * (CYTOSOLIC\_PER\_CRY\_PROTEIN) - Kpc1 * (CYTOSOLIC\_PER\_CRY\_PROTEIN) - Kpcp{*}(CYTOSOLIC\_PER\_CRY\_PROTEIN) + Kpcpc{*}(PHOS\_CYTOSOLIC\_PER\_CRY\_PROTEIN) - dpcc * (CYTOSOLIC\_PER\_CRY\_PROTEIN) \quad (11)$$

• The next six ordinary differential equations of this model represent the reversible activities of the phophorylated PER, CRY, BMAL1, REV-ERB, ROR protein and PER-CRY protein complex in the cytoplasm denoted by $P_{pc},\ C_{pc},\ B_{pc},\ Re_{pc}$ , $Ro_{pc}$ & $PC_p$ respectively.

$$\frac{dP_{pc}}{dt} = Kpc * (CYTOSOLIC\_PER\_PROTEIN) - Kppc * (PHOS\_CYTOSOLIC\_PER\_PROTEIN) - dppc * (PHOS\_CYTOSOLIC\_PER\_PROTEIN) \quad (12)$$

$$\frac{dC_{pc}}{dt} = Kcc * (CYTOSOLIC\_CRY\_PROTEIN) - Kcpc * (PHOS\_CYTOSOLIC\_CRY\_PROTEIN) - dcpc * (PHOS\_CYTOSOLIC\_CRY\_PROTEIN) \quad (13)$$

$$\frac{dB_{pc}}{dt} = Kbc * (CYTOSOLIC\_BMAL1\_PROTEIN) - Kbpc{*}(PHOS\_CYTOSOLIC\_BMAL1\_PROTEIN) - dbc * (PHOS\_CYTOSOLIC\_BMAL1\_PROTEIN) \quad (14)$$

$$\frac{dRo_{pc}}{dt} = Krc * ((CYTOSOLIC\_ROR\_PROTEIN)) - Krpc * (PHOS\_CYTOSOLIC\_ROR\_PROTEIN) - drc * (PHOS\_CYTOSOLIC\_ROR\_PROTEIN) \quad (15)$$

$$\frac{dRe_{pc}}{dt} = Krec * ((CYTOSOLIC\_REV\_ERB\_PROTEIN)) - Krepc*(PHOS\_CYTOSOLIC\_REV\_ERB\_PROTEIN) - drec * (PHOS\_CYTOSOLIC\_REV\_ERB\_PROTEIN) \quad (16)$$

$$\frac{dPC_{pc}}{dt} = Kpcp * (CYTOSOLIC\_PER\_CRY\_PROTEIN) - Kpcpc*(PHOS\_CYTOSOLIC\_PER\_CRY\_PROTEIN) - dpcc * (PHOS\_CYTOSOLIC\_PER\_CRY\_PROTEIN) \quad (17)$$

• The last six differential equations of this model represent the activites of BMAL1, ROR, REV-ERB, CLOCK-BMAL, PER-CRY, and PER-CRY/CLOCK-BMAL protein in the nucleus denoted by $B_n$ ,$Ro_n$, $Re_n$, $CB_n$, $PC_n$, $PC/CB_n$, respectively.

$$\frac{dB_n}{dt} = Kbcc * (CYTOSOLIC\_BMAL1\_PROTEIN) - Kclbn * (NUCLEAR\_BMAL1\_PROTEIN) - dbn. * (NUCLEAR\_BMAL1\_PROTEIN) \quad (18)$$

$$\frac{dRo_n}{dt} = Krcc * (CYTOSOLIC\_ROR\_PROTEIN) - Krn * (NUCLEAR\_ROR\_PROTEIN) - drn * (NUCLEAR\_ROR\_PROTEIN) \quad (19)$$

$$\frac{dRe_n}{dt} = Krecc * (CYTOSOLIC\_REV\_ERB\_PROTEIN) - Kren * (NUCLEAR\_REV\_ERB\_PROTEIN) - dren * (NUCLEAR\_REV\_ERB\_PROTEIN); \quad (20)$$

$$\frac{dCB_n}{dt} = Kclbn * (NUCLEAR\_BMAL1\_PROTEIN) - Kcbpc * (NUCLEAR\_CLOCK\_BMAL1\_PROTEIN) * (NUCLEAR\_PER\_CRY\_PROTEIN) + Kdcbpc * (NUCLEAR\_CLOCK\_BMAL1\_PER\_CRY\_PROTEIN) - dclbn*(NUCLEAR\_CLOCK\_BMAL1\_PROTEIN) + dpcn*(NUCLEAR\_CLOCK\_BMAL1\_PER\_CRY\_PROTEIN) \quad (21)$$

$$\frac{dPC_n}{dt} = Kpcc * (CYTOSOLIC\_PER\_CRY\_PROTEIN) - Kcbpc * (NUCLEAR\_CLOCK\_BMAL1\_PROTEIN) * (NUCLEAR\_PER\_CRY\_PROTEIN) + Kdcbpc * (NUCLEAR\_CLOCK\_BMAL1\_PER\_CRY\_PROTEIN) - dpcn*(NUCLEAR\_PER\_CRY\_PROTEIN) + dclbn*(NUCLEAR\_CLOCK\_BMAL1\_PER\_CRY\_PROTEIN) \quad (22)$$

$$\frac{dPC/CB_n}{dt} = Kcbpc * (NUCLEAR\_CLOCK\_BMAL1\_PROTEIN) * (NUCLEAR\_PER\_CRY\_PROTEIN) - Kdcbpc * (NUCLEAR\_CLOCK\_BMAL1\_PER\_CRY\_PROTEIN) - dclbn * (NUCLEAR\_CLOCK\_BMAL1\_PER\_CRY\_PROTEIN) - dpcn * (NUCLEAR\_CLOCK\_BMAL1\_PER\_CRY\_PROTEIN) \quad (23)$$

**Model 2: Model with communication between the cellular oscillators**

To incorporate the synchronicity of cells in the second model, we assume a global coupling among the cells through the cell to cell communication between $N$ number of cells. Using the parameter $K_f$ to describe the communication strength, we calculated the parameter $F$, which is the mean communication signal among $N$ number of cells in the system using compucel3d which was used in the transcription equation of each the five coding genes to relay the coupling mechanism in the model. This approach was initially used in a three variable model by *Gonze D et al, 2005*. We assume the cell to cell communication between the cells leading to synchronization is mediated by a coupling protein. Previous studies has assumed the Transforming growth factor beta (TGF-$\beta$) to be the coupling factor responsible for the cell to cell communication and the synchronization of the cells to produce a sustained oscillation with synchronized phase and period (Finger A et al, 2021).

• The evolution equation for a single synchronization factor ($M$, which will later be used to calculate the mean field $F$) in the cellular medium by a coupling protein is given as:

$$\frac{dM}{dt} = prt * X_i - \frac{vc * M}{dc + M} \tag{24}$$

Where $i = 1...5$ represents each of the observable genes used in the model. $X_1$ for *Per*, $X_2$ for *Cry*, $X_3$ for*Bmal1*, $X_4$ for *Ror*, and $X_5$ for *Rev-erb*. The mean field $F$ was then calculated as:

$$F = \frac{1}{N} * \sum_{i=1}^{N} M_i \tag{25}$$

• The first five system of ordinary differential equation of this model represents the transcription of *Per*, *Cry*, *Bmal1*, *Ror* & *Rev-erb* genes into their respective mRNA's denoted by $M_P$, $M_C$, $M_B$, $M_{Ro}$ & $M_{Re}$ respectively by the nuclear $CLOCK\_BMAL$ protein ($CB_n$), $PER\_CRY$ protein ($PC_n$), $ROR$ protein ($Ro_n$) and $REV\_ERB$ protein ($Re_n$).

$$\frac{dM_P}{dt} = light * \frac{V_{sp} * (CLOCK\_BMAL1)^n}{Kip^n + (CLOCK\_BMAL1 * PER\_CRY)^m + (CLOCK\_BMAL1)^n} + \frac{Vdp * K_f * F}{K_c + K_f * F} - d_p * M_P \tag{26}$$

$$\frac{dM_C}{dt} = light * \frac{light * V_{sc} * (CLOCK\_BMAL1)^o}{Kic^o + (CLOCK\_BMAL1 * PER\_CRY)^p + (CLOCK\_BMAL1)^o} + \frac{Vdp * K_f * F}{K_c + K_f * F} - d_c * M_C \tag{27}$$

$$\frac{dM_B}{dt} = \frac{V_{sb} * (ROR)^p}{Kib^p + (ROR * REV\_ERB)^q + (ROR)^p} + \frac{Vdp * K_f * F}{K_c + K_f * F} - d_b * M_B \tag{28}$$

$$\frac{dM_R}{dt} = \frac{V_{sr} * (CLOCK\_BMAL1)^r}{Kir^r + (CLOCK\_BMAL1 * PER\_CRY)^s + (CLOCK\_BMAL1)^r} + \frac{Vdp * K_f * F}{K_c + K_f * F} - d_r * M_R \tag{29}$$

$$\frac{dM_{Re}}{dt} = \frac{V_{sre} * (CLOCK\_BMAL1)^t}{Kire^t + (CLOCK\_BMAL1 * PER\_CRY)^u + (CLOCK\_BMAL1)^t} + \frac{Vdp * K_f * F}{K_c + K_f * F} - d_{re} * M_{Re} \tag{30}$$

• The next six system of ordinary differential equations of this model represents the translation and reversible activities of the unphophorylated PER, CRY, BMAL1, REV-ERB, ROR protein and PER-CRY protein complex in the cytoplasm denoted by $P_c$, $C_c$, $B_c$, $Re_c$ , $Ro_c$ & $PC_c$ respectively.

$$\begin{aligned}\frac{dP_c}{dt} &= k * Per\_mRNA + Kpc1*(CYTOSOLIC\_PER\_CRY\_PROTEIN) - Kpco*(CYTOSOLIC\_PER\_PROTEIN) \\ &* (CYTOSOLIC\_CRY\_PROTEIN) - Kpc * ((CYTOSOLIC\_PER\_PROTEIN)) + \\ &Kppc * (PHOS\_CYTOSOLIC\_PER\_PROTEIN) - dpc * (CYTOSOLIC\_PER\_PROTEIN)\end{aligned} \tag{31}$$

$$\begin{aligned}\frac{dC_c}{dt} &= k1 * Cry\_mRNA + Kpc1*(CYTOSOLIC\_PER\_CRY\_PROTEIN) - Kpco*(CYTOSOLIC\_PER\_PROTEIN) \\ &* (CYTOSOLIC\_CRY\_PROTEIN) - Kcc * ((CYTOSOLIC\_CRY\_PROTEIN)) \\ &+ Kcpc * (PHOS\_CYTOSOLIC\_CRY\_PROTEIN) - dcc * (CYTOSOLIC\_CRY\_PROTEIN)\end{aligned} \tag{32}$$

$$\begin{aligned}\frac{dB_c}{dt} &= k2 * Bmal1\_mRNA - Kbcc*(CYTOSOLIC\_BMAL1\_PROTEIN) - Kbc*((CYTOSOLIC\_BMAL1\_PROTEIN)) \\ &+ Kbpc * (PHOS\_CYTOSOLIC\_BMAL1\_PROTEIN) - dbc * (CYTOSOLIC\_BMAL1\_PROTEIN)\end{aligned} \tag{33}$$

$$\frac{dRo_c}{dt} = k3 * Ror\_mRNA - Krcc{*}(CYTOSOLIC\_ROR\_PROTEIN) - Krc{*}((CYTOSOLIC\_ROR\_PROTEIN)) + Krpc * (PHOS\_CYTOSOLIC\_ROR\_PROTEIN) - drc * (CYTOSOLIC\_ROR\_PROTEIN) \quad (34)$$

$$\frac{dRe_c}{dt} = k4 * Rev\ _erb\ _mRNA - Krecc * (CYTOSOLIC\_REV\_ERB\_PROTEIN) - Krec * ((CYTOSOLIC\_REV\_ERB\_PROTEIN)) + Krepc * (PHOS\_CYTOSOLIC\_REV\_ERB\_PROTEIN) - drec * (CYTOSOLIC\_REV\_ERB\_PROTEIN) \quad (35)$$

$$\frac{dPC_c}{dt} = Kpco * ((CYTOSOLIC\_PER\_PROTEIN) * (CYTOSOLIC\_CRY\_PROTEIN)) - Kpcc * (CYTOSOLIC\_PER\_CRY\_PROTEIN) - Kpc1 * (CYTOSOLIC\_PER\_CRY\_PROTEIN) - Kpcp{*}(CYTOSOLIC\_PER\_CRY\_PROTEIN) + Kpcpc{*}(PHOS\_CYTOSOLIC\_PER\_CRY\_PROTEIN) - dpcc * (CYTOSOLIC\_PER\_CRY\_PROTEIN) \quad (36)$$

• The next six system of ordinary differential equations of this model represents the reversible activities of the phophorylated PER, CRY, BMAL1, REV-ERB, ROR protein and PER-CRY protein complex in the cytoplasm denoted by $P_{pc}$, $C_{pc}$, $B_{pc}$, $Re_{pc}$ , $Ro_{pc}$ & $PC_{pc}$ respectively.

$$\frac{dP_{pc}}{dt} = Kpc * (CYTOSOLIC\_PER\_PROTEIN) - Kppc * (PHOS\_CYTOSOLIC\_PER\_PROTEIN) - dppc * (PHOS\_CYTOSOLIC\_PER\_PROTEIN) \quad (37)$$

$$\frac{dC_{pc}}{dt} = Kcc * (CYTOSOLIC\_CRY\_PROTEIN) - Kcpc * (PHOS\_CYTOSOLIC\_CRY\_PROTEIN) - dcpc * (PHOS\_CYTOSOLIC\_CRY\_PROTEIN) \quad (38)$$

$$\frac{dB_{pc}}{dt} = Kbc * (CYTOSOLIC\_BMAL1\_PROTEIN) - Kbpc{*}(PHOS\_CYTOSOLIC\_BMAL1\_PROTEIN) - dbc * (PHOS\_CYTOSOLIC\_BMAL1\_PROTEIN) \quad (39)$$

$$\frac{dRo_{pc}}{dt} = Krc * ((CYTOSOLIC\_ROR\_PROTEIN)) - Krpc * (PHOS\_CYTOSOLIC\_ROR\_PROTEIN) - drc * (PHOS\_CYTOSOLIC\_ROR\_PROTEIN) \quad (40)$$

$$\frac{dRe_{pc}}{dt} = Krec * ((CYTOSOLIC\_REV\_ERB\_PROTEIN)) - Krepc{*}(PHOS\_CYTOSOLIC\_REV\_ERB\_PROTEIN) - drec * (PHOS\_CYTOSOLIC\_REV\_ERB\_PROTEIN) \quad (41)$$

$$\frac{dPC_{pc}}{dt} = Kpcp * (CYTOSOLIC\_PER\_CRY\_PROTEIN) - Kpcpc{*}(PHOS\_CYTOSOLIC\_PER\_CRY\_PROTEIN) - dpcc * (PHOS\_CYTOSOLIC\_PER\_CRY\_PROTEIN) \quad (42)$$

• The last six differential equations of this model represents the activites of BMAL1, ROR, REV-ERB, CLOCK-BMAL, PER-CRY, and PER-CRY/CLOCK-BMAL protein in the nucleus denoted by $B_n$ , $Ro_n$, $Re_n$, $CB_n$, $PC_n$, $PC/CB_n$, respectively.

$$\frac{dB_n}{dt} = Kbcc \, * \, (CYTOSOLIC\_BMAL1\_PROTEIN) \, - \, Kclbn * (NUCLEAR\_BMAL1\_PROTEIN) - dbn. * (NUCLEAR\_BMAL1\_PROTEIN) \quad (43)$$

$$\frac{dRo_n}{dt} = Krcc \, * \, (CYTOSOLIC\_ROR\_PROTEIN) \, - \, Krn * (NUCLEAR\_ROR\_PROTEIN) - drn * (NUCLEAR\_ROR\_PROTEIN) \quad (44)$$

$$\frac{dRe_n}{dt} = Krecc \, * \, (CYTOSOLIC\_REV\_ERB\_PROTEIN) \, - \, Kren * (NUCLEAR\_REV\_ERB\_PROTEIN) - dren * (NUCLEAR\_REV\_ERB\_PROTEIN); \quad (45)$$

$$\frac{dCB_n}{dt} = Kclbn \, * \, (NUCLEAR\_BMAL1\_PROTEIN) - Kcbpc * (NUCLEAR\_CLOCK\_BMAL1\_PROTEIN) * (NUCLEAR\_PER\_CRY\_PROTEIN) + \, Kdcbpc * (NUCLEAR\_CLOCK\_BMAL1\_PER\_CRY\_PROTEIN) - dclbn*(NUCLEAR\_CLOCK\_BMAL1\_PROTEIN) + dpcn*(NUCLEAR\_CLOCK\_BMAL1\_PER\_CRY\_PROTEIN) \quad (46)$$

$$\frac{dPC_n}{dt} = Kpcc \, * \, (CYTOSOLIC\_PER\_CRY\_PROTEIN) - Kcbpc * (NUCLEAR\_CLOCK\_BMAL1\_PROTEIN) * (NUCLEAR\_PER\_CRY\_PROTEIN) + \, Kdcbpc * (NUCLEAR\_CLOCK\_BMAL1\_PER\_CRY\_PROTEIN) - dpcn*(NUCLEAR\_PER\_CRY\_PROTEIN) + dclbn*(NUCLEAR\_CLOCK\_BMAL1\_PER\_CRY\_PROTEIN) \quad (47)$$

$$\frac{dPC/CB_n}{dt} = Kcbpc \, * \, (NUCLEAR\_CLOCK\_BMAL1\_PROTEIN) * (NUCLEAR\_PER\_CRY\_PROTEIN) - \, Kdcbpc * (NUCLEAR\_CLOCK\_BMAL1\_PER\_CRY\_PROTEIN) - \, dclbn * (NUCLEAR\_CLOCK\_BMAL1\_PER\_CRY\_PROTEIN) - dpcn * (NUCLEAR\_CLOCK\_BMAL1\_PER\_CRY\_PROTEIN) \quad (48)$$

| Parameter | Definition | Set 1 | Set 2 | Set 2 |
|---|---|---|---|---|
| Vsp | Maximum rate of Per synthesis | 2.2 | 1.7 | 3.0 |
| Kip | Activation constant for enhancement of Per by nuclear CLOCK/BMAL | 0.24 | 0.302 | 0.28 |
| d_p | Maximum rate of Per degradation | 0.34 | 0.375 | 0.429 |
| Vsc | Maximum rate of Cry synthesis | 2.3 | 1.6 | 2.8 |
| Kic | Activation constant for enhancement of Cry by nuclear CLOCK/BMAL | 0.262 | 0.32 | 0.17 |
| d_c | Maximum rate of Cry degradation | 0.324 | 39 | 0.44 |
| Vsb | Maximum rate of Bmal1 synthesis | 2.32 | 1.2 | 3.4 |
| Kib | Activation constant for enhancement of Cry by nuclear Ror | 0.13 | 0.16 | 0.16 |
| d_b | Maximum rate of Bmal1 degradation | 0.46 | 0.43 | 0.49 |
| Vsr | Maximum rate of Ror synthesis | 2.04 | 1.6 | 3.1 |
| Kir | Activation constant for enhancement of Ror by nuclear CLOCK/BMAL | 0.27 | 0.39 | 0.3 |
| d_r | Maximum rate of Ror degradation | 0.372 | 0.34 | 0.27 |
| Vsre | Maximum rate of Rev-erb synthesis | 2.15 | 1.8 | 2.9 |
| Kic | Activation constant for enhancement of Rev-erb by nuclear CLOCK/BMAL | 0.24 | 0.37 | 0.25 |
| d_re | Maximum rate of Rev-erb degradation | 0.382 | 0.32 | 0.38 |
| k | Maximum rate of Per mRNA translation | 0.408 | 0.453 | 0.412 |
| Kpc1 | Dissociation rate of PER-CRY protein to PER protein | 0.362 | 0.383 | 0.34 |
| Kpco | Association rate of PER & CRY protein to PER-CRY protein | 0.3 | 0.484 | 0.31 |
| Kpc | Rate of PER phosphorylation | 0.304 | 0.204 | 0.33 |
| Kppc | Rate of PER de-phosphorylation | 0.39 | 0.306 | 0.32 |
| dpc | Degradation Rate of PER Protein | 0.23 | 0.024 | 0.10 |
| k1 | Maximum rate of Cry mRNA translation | 0.39 | 0.53 | 0.42 |
| Kcc | Rate of CRY phosphorylation | 0.365 | 0.242 | 0.4 |
| Kcpc | Rate of CRY de-phosphorylation | 0.306 | 0.31 | 0.35 |
| dpc | Degradation Rate of PER Protein | 0.18 | 0.023 | 0.08 |
| k2 | Maximum rate of Bmal1 mRNA translation | 0.47 | 0.34 | 0.53 |
| Kbcc | Rate of cytosol BMAL Protein transported into the nucleus | 0.34 | 0.342 | 0.48 |
| Kbc | Rate of BMAL phosphorylation | 0.30 | 0.28 | 0.3 |
| Kbpc | Rate of BMAL de-phosphorylation | 0.38 | 0.39 | 0.25 |
| dbc | Degradation Rate of BMAL Protein | 0.07 | 0.05 | 0.07 |
| k3 | Maximum rate of Ror mRNA translation | 0.42 | 0.44 | 0.5 |
| Krcc | Rate of cytosol ROR Protein transported into the nucleus | 0.4 | 0.35 | 0.504 |
| Krc | Rate of ROR phosphorylation | 0.33 | 0.36 | 0.4 |
| Krpc | Rate of ROR de-phosphorylation | 0.2 | 0.40 | 0.16 |
| drc | Degradation Rate of ROR Protein | 0.05 | 0.05 | 0.05 |
| k4 | Maximum rate of Rev-erb mRNA translation | 0.42 | 0.36 | 0.4 |
| Krecc | Rate of cytosol REV-ERB Protein transported into the nucleus | 0.4 | 0.304 | 0.48 |
| Krec | Rate of REV-ERB phosphorylation | 0.33 | 0.36 | 0.36 |
| Krepc | Rate of REV-ERB de-phosphorylation | 0.2 | 0.34 | 0.33 |
| drec | Degradation Rate of REV-ERB Protein | 0.05 | 0.06 | 0.05 |
| Kpcc | Rate of cytosol PER-CRY Protein transported into the nucleus | 0.37 | 0.345 | 0.4 |
| Kpcp | Rate of PER-CRY phosphorylation | 0.25 | 0.24 | 0.28 |
| Kpcpc | Rate of PER-CRY de-phosphorylation | 0.36 | 0.36 | 0.39 |
| dpcc | Degradation Rate of PER-CRY Protein | 0.017 | 0.017 | 0.017 |
| dppc | Degradation Rate of Phosphorylated PER Protein | 0.023 | 0.023 | 0.23 |
| dcpc | Degradation Rate of Phosphorylated CRY Protein | 0.020 | 0.019 | 0.020 |
| dbpc | Degradation Rate of Phosphorylated BMAL Protein | 0.013 | 0.013 | 0.015 |
| drpc | Degradation Rate of Phosphorylated ROR Protein | 0.02 | 0.02 | 0.02 |
| drepc | Degradation Rate of Phosphorylated REV-ERB Protein | 0.023 | 0.23 | 0.02 |
| dpcpc | Degradation Rate of Phosphorylated PER-CRY Protein | 0.025 | 0.025 | 0.03 |
| Kclbn | Association rate of CLOCK and BMAL protein to form CLOCK-BMAL protein | 0.37 | 0.32 | 0.47 |
| dbn | Degradation Rate of nuclear BMAL Protein | 0.09 | 0.03 | 0.05 |
| Krn | Association rate of ROR and REV-ERB protein | 0.35 | 0.31 | 0.25 |
| drn | Degradation Rate of nuclear ROR Protein | 0.07 | 0.02 | 0.07 |
| Kren | Association rate of REV-ERB and ROR protein | 0.35 | 0.31 | 0.35 |
| drn | Degradation Rate of nuclear REV-ERB Protein | 0.06 | 0.02 | 0.06 |
| Kcbpc | Association rate of CLOCK-BMAL & PER-CRY proteins in the nucleus | 0.49 | 0.25 | 0.4 |
| dclbn | Degradation Rate of nuclear CLOCK-BMAL Protein | 0.30 | 0.15 | 0.30 |
| dpcn | Degradation Rate of nuclear PER-CRY Protein | 0.15 | 0.075 | 15 |
| prt | synchronization protein concentration | 1.34 | 0.945 | 1.28 |
| vc | Maximum rate of synchronization factor synthesis | 1.734 | 1.945 | 1.58 |
| dc | Activation constant for enhancement of synchronization factor synthesisL | 0.134 | 0.145 | 0.28 |
